# Cooperative breeding and the emergence of multilevel societies in birds

**DOI:** 10.1101/2021.10.04.462904

**Authors:** Ettore Camerlenghi, Alexandra McQueen, Kaspar Delhey, Carly N. Cook, Sjouke A. Kingma, Damien R. Farine, Anne Peters

**Author notes:** **Corresponding author:** Ettore Camerlenghi, /.

## Abstract

Multilevel societies (MLSs), where social levels are hierarchically nested within each other, are considered one of the most complex forms of animal societies. Although thought to mainly occur in mammals, it is suggested that MLSs could be under-detected in birds. Here we propose that the emergence of MLSs could be common in cooperatively breeding birds, as both systems are favoured by similar ecological and social drivers. We first investigate this proposition by systematically comparing evidence for multilevel social structure in cooperative and non-cooperative birds in Australia and New Zealand, global hotspots for cooperative breeding. We then analyse non-breeding social networks of cooperatively breeding superb fairy-wrens (*Malurus cyaneus*) to reveal their structured multilevel society, with three hierarchical social levels that are stable across years. Our results confirm recent predictions that MLSs are likely to be widespread in birds and suggest that these societies could be particularly common in cooperatively breeding birds.

## INTRODUCTION

Multilevel societies (MLSs) represent social structures that comprise two or more social levels nested within each other (e.g., family units or breeding pairs that form bands, and bands that form clans), and where units at each level (other than the upper-most) are socially cohesive (Grueter et al. 2020). MLSs were first described in Hamadryas baboons (*Papio hamadryas*) (Kummer 1968) and subsequently in other primates (Kirkpatrick and Grueter 2010; Snyder-Mackler et al. 2012), with a number of two- and three-level societies having subsequently been described in other mammalian taxa, such as African elephants (*Loxodonta africana*) (Wittemyer et al. 2005), sperm whales (*Physeter macrocephalus*) (Whitehead 2012) and killer whales (*Orcinus orca*) (Tavares et al. 2016). The vast majority of human societies also show a multi-tiered structure to their social system, particularly hunter-gatherer societies that often consist of families that are part of local bands, which in turn form higher-level tribes (Chapais 2011; Migliano et al. 2017; Grueter 2017; Grueter et al. 2020). The need for individuals in MLSs to track and maintain highly differentiated relationships across various levels—potentially spanning a large number of conspecifics—makes multilevel societies one of the most complex forms of social structures in vertebrates (Swedell and Plummer 2012, but see Papageorgiou et al. 2019). As a result, studies of MLSs are considered particularly important for understanding the evolution of animal sociality and to shed light on the evolution of human societies (Grueter et al. 2017). Nevertheless, we still have limited understanding of the drivers and functions of this type of social structure.

Multilevel societies are hypothesised to arise as a means of giving individuals opportunities to adopt flexible social responses to environmental variability without compromising the integrity and composition of the social units (Kummer 1968). Because MLSs allow groups to increase or decrease in size as conditions become more or less unpredictable, MLSs are thought to provide members with a distinct advantage over species in which group size is fixed (Grueter 2017). For example, Hamadryas baboons predominantly live in one-male multi-female units, but at sleeping cliffs, water holes and in seasons where food is abundant, family groups coalesce into clans, then bands and troops to collectively defend resources and reduce predation risk (Schreier and Swedell 2009). By being able to split back into family units, baboons can find sufficient food when the resources are limited and/or dispersed (Schreier and Swedell 2012). While dynamically forming and disbanding social levels can be beneficial by allowing animals to solve problems that fixed social units cannot (Grueter 2017), it remains unclear why some species form several nested social units whereas in others, units remain agonistic to one-another and, in many cases, strictly territorial (Grueter et al. 2020). Unravelling drivers of MLS is therefore important because it would allow inferences to be made about the ecological, social and genetic factors that underlie the evolution of sociality and cooperation between social units.

Although research initially focused predominately on primates, there is now ample evidence that MLSs are more common than previously thought (Whitehead 2012; Wittemyer et al. 2005; Rubenstein and Hack 2004; Bigg et al.1990; Papageorgiou and Farine 2020). Recently, it has also been shown that a bird (the vulturine guineafowl *Acryllium vulturinum*) exhibits a multilevel social structure (Papageorgiou et al. 2019). This species lives in the open woodlands of East Africa, and experiences similar ecological conditions to those of elephants and hamadryas baboons (Del Hoyo et al.1994), which suggests that specific ecological factors, rather than cognitive capacity, might play a role in driving the formation of MLS (Papageorgiou et al. 2019). Thus, focusing on population-level social behaviour across species experiencing different ecological conditions could provide new insights into some of the drivers leading to the evolution of multilevel societies (Papageorgiou and Farine 2020).

A number of examples of nested societies are known to exist in birds, but these have only recently started to be considered within the framework of MLSs (Papageorgiou and Farine 2020). For example, bell miners (*Manorina melanophyris*) (Clarke and Fitz-Gerald 1994; Painter et al. 2000), white-fronted bee-eaters (*Merops bullockoides*) (Hegner et al.1982) and buff-rumped thornbills (*Acanthiza reguloides*) (Bell and Ford 1986) are known to live in societies structured into breeding groups nested within clans. If these social levels were shown to be consistent and stable, this would result in these bird societies meeting the criteria for being identified as MLSs (Grueter et al. 2020 b). One aspect that these species also have in common is that they all breed cooperatively, raising the possibility that cooperative breeding might facilitate, or be driven by similar factors to, the formation of MLSs. A few arguments support this proposition. In mammals, MLS are commonly found in kin-structured societies (Patzelt et al. 2014; Stäedele et al. 2015). Cooperatively breeding species similarly live in family-units for consecutive seasons, where some individuals stay in their natal territory to help breeders defend a territory and raise young, yielding direct and indirect benefits (Cockburn 1998; Kingma et al. 2014; Koenig and Dickinson 2016; Downing et al. 2020). While the presence of family units in-and-of-itself should not predispose cooperative breeders to form MLSs (i.e. this represents the base level, which equally applies to pair-living species), cooperative breeding is often linked to strong kin structure extending beyond the territory because habitat saturation often limits dispersal distances (Painter et al. 2000).

As a result of philopatry and delayed dispersal, populations of cooperative breeders generally show high social viscosity (Hamilton 1964 b), a spatial pattern of relatedness where related individuals remain in close proximity (Wolf and Trillmich 2008). This gives rise to both direct and indirect benefits if grouping with related individuals beyond the core group, for example during the non-breeding season, is beneficial. Such benefits could be most important during harsh conditions, when there is greater need for information transfer about food availability (Aplin et al. 2012), or group vigilance and defence due to high predation risk (Kenward 1978). Environmental harshness (or unpredictability) is suggested to be linked to cooperative breeding (Jetz and Rubenstein 2011), in line with the prediction that MLS enable social groups to deal with environmental variability (Städele et al. 2016; Schreier and Swedell 2009). Thus, the factors that are thought to favour cooperative breeding appear to be the same as those thought to underlie the formation of MLS.

Here, we propose that cooperative breeding could facilitate the emergence of multilevel societies, which, in accordance with Grueter et al. (2020), we define as social structure exhibiting (i) consistency of individual membership in each level over time, and (ii) spatio-temporal cohesion of the core and upper levels. To test our proposition, we first compare the evidence in the literature suggestive of multilevel social structure in cooperatively breeding versus non-cooperatively breeding birds. We focus our search on Australian and New Zealand avifauna because of the relatively high-quality information available on the social structure of these species, and the preponderance of cooperative breeding in this geographical region (Griesser et al. 2017). Second, we use empirical data collected from a cooperatively breeding species identified as putatively displaying MLS to formally test for the presence of a multilevel social structure using modern social analytical approaches.

## METHODS

### Comparative analysis of cooperatively breeding birds

We performed a systematic literature search for evidence suggestive of the existence of multilevel societies across the Australian and New Zealand bird families that are known to include at least one cooperative breeding species [n = 17 families; 43 cooperatively breeding species and 70 non-cooperative species (Cornwallis et al. 2017)]. Following Cockburn (2006) and Cornwallis et al. (2017), we define cooperative breeding as a breeding system in which at least 10% of young are retained on their natal territory and provide care to the offspring of a breeding pair.

We collected the information available on social structure from the detailed species accounts of Handbook of Australian, New Zealand and Antarctic birds (HANZAB) (Marchant et al. 2006), particularly the sections on social organisation and behaviour. These represent a comprehensive summary of the knowledge of the avifauna of Australia, New Zealand and Antarctica until 2006. We then updated this information by searching for studies published after 2000. We conducted a step-wise structured search in Web of Science using the following search strings: 1) “common name of the species” AND (social* OR winter OR aggregation OR congregation); and 2) “scientific name of the species” AND (social* OR winter OR aggregation OR congregation). For the species for which no records or very few records were found, we repeated the search with 3) “common name of the species*” (including various spellings and * for the plural) and OR “scientific name of the species”. The title and abstract of each study were screened to determine if it met the inclusion criteria: information about sociality, social organisation, social structure, or demography of the species. Studies that met at least one of these criteria (n=100 out of 965 articles) were read in full, recording the information on the social organisation or social structure of the species during both breeding and non-breeding season.

Species for which any sources suggested they form at least one higher social unit than the breeding group/pair were classified as potential MLS. Our criteria included: repeated aggregations of breeding groups/family groups/breeding pairs; formation of stable foraging flocks, clans, bands; supergroups, which were reported to be stable but also were observed to separate periodically into the original breeding pairs/groups. Species reported to form large and loose flocks or aggregations, or to defend a territory year-round, were not considered to exhibit MLS.

To assess whether cooperative breeding predicts the presumed presence of multilevel societies, we used phylogenetic logistic regression with the function phyloglm in the R package phylolm (Ho and Ane 2014) with MLS as binary response, and cooperative or not as explanatory variable. The model also included the length of text (cm) devoted to the social organisation of the species in the HANZAB as a covariate to account for the possibility that species for which more information is available may be more likely to be considered to have MLS. Numerical covariate (length of text) was scaled to zero-mean and unit-variance. We incorporated phylogenetic uncertainty into the error estimates of each parameter by running the model across 100 phylogenies (obtained from www.birdtree.org), using the Hackett backbone (Jetz et al. 2012), and summarised parameters using Rubin’s rules (Nakagawa and De Villemereuil 2019). We computed relative efficiency for each model parameter to determine the efficacy of accounting for phylogenetic uncertainty. This parameter varies between 0 and 1, and quantifies the efficiency achieved with the number of phylogenies used relative to the theoretical (maximal) efficiency obtained by using an infinite number of phylogenies. Values exceeded the recommended value of 0.99 (Nakagawa and De Villemereuil 2019) for all parameters, which suggests that 100 phylogenies adequately capture phylogenetic uncertainty.

### Field case study

#### Field study system

We studied the social behaviour and movement of a population of superb fairy-wrens *Malurus cyaneus*, a small (9–12 g) facultative cooperatively breeding songbird. We observed the social structure of the population across two years, with a particular focus on the behaviour of individuals during the non-breeding season. Fairy-wrens form stable pairs or groups during the breeding season which actively defend their exclusive breeding territory from conspecific intruders (Rowley 1964; Cooney and Cockburn 1995). However, during the rest of the year individuals from multiple breeding territories can aggregate to form larger groups (Fallow and Magrath 2009). We quantified this pattern at Lysterfield Park reserve, located near Melbourne, Australia (−37° 56’ 56.40” S, 145° 17’ 45.60” E) on the traditional land of Bunurong Boon Wurrung and Wurundjeri Woi Wurrung peoples of the Eastern Kulin Nation. The study area is comprised of open woodland, including areas with dense shrubs and open grassland. Almost all (98%) individuals in the study (n=198) were colour banded with a unique combination of two metal rings, and two plastic coloured rings. All birds were sexed based on morphological traits (males develop a black bill and male breeding plumage in their first Spring; females and juveniles are indistinguishable) with the exception of 13 individuals, which disappeared before exhibiting male characteristics. Of these, nine were sexed as females by PCR (following methods in Eastwood et al. 2018). For the other four, no blood sample was collected; these individuals were also presumed females.

#### Field Observations

We observed all individuals throughout 2018, 2019 and 2020 as part of a long-term study. During the breeding seasons (September-January/February) we identified 19 territories and calculated their total area, based on movement of the birds, using GPS. The identity of all individuals in the breeding groups was recorded during weekly observations. Reproductive attempts were followed and the offspring were banded. Over both breeding seasons (2018-2019 and 2019-2020) eight of the 19 breeding groups had one or two helpers, and the average 2018-2019 breeding group size (without counting the offspring of the same breeding season) was 2.7 individuals per group or 3.2 when considering new offspring. During breeding season 2019-2020, the average breeding group size was 2.5 individuals or 5.8 individuals when considering new offspring.

During the months before and after each breeding season (March-April and September), all individuals were sighted weekly by systematically searching all territories and the broader surrounding area. Throughout the non-breeding season (May-August), groups coalesce into larger roaming flocks, which we defined as temporary aggregations of individuals in the same place at the same time (following Shizuka et al. 2014). During these months, we systematically searched for flocks. Whenever a flock was encountered, we followed all individuals in the flock for a minimum of 30 minutes. As in previous studies on related species (e.g. Farine & Milburn 2013), flock membership during observations was unambiguous, with flocks of individuals moving together and being separated from other flocks by a considerable distance. While following flocks, we recorded the movement of the flock at intervals of five minutes with a handheld GPS (for later determination of home range size). We assigned each flock a unique identifier code and recorded the identity of all individuals in the flock using their unique combination of colour rings, along with the time and GPS coordinates of the observation.

#### Constructing social networks and identifying social units

We used the *asnipe* package (Farine 2013) in R (R Core Team, 2020) to construct a group by individual matrix for each non-breeding season, with every individual in the population as a column and every unique flock as a row, with cell values 0 and 1 corresponding to whether the individual was absent or present in a given flock. From these matrices, we calculated a social network for the population using the Simple Ratio Index (Hoppitt and Farine 2018). We omitted from the networks 85 colour banded individuals that were observed fewer than five times per season and which disappeared from the study area during the first month of the data collection, as these would have generated excessive uncertainty in the resulting network (Davis et al. 2018). These transient individuals appeared to be mostly dispersing juvenile females, as they were all in female/juvenile plumage and generally observed only once. We constructed two networks, representing the social structure of our study population during May-August in 2019 and 2020.

To identify social units across different levels, we first plotted a histogram of association strengths, and noted that this distribution was multimodal. We first defined non-breeding groups from the upper-most mode, comprising pairs of individuals that were observed together more than 75% of the time. Then, following Papageorgiou et al (2019), we identified the upper-level communities in the networks, and their composition, by running the ‘fastgreedy’ statistical community-detection algorithm in the R package igraph (Csardi and Nepusz 2006). We tested the robustness of the composition of the non-breeding groups and communities by running the algorithm described by Shizuka & Farine (2016). When testing the robustness of the communities’ composition, this algorithm bootstraps the observed group by individual matrix, constructs a social network from each bootstrapped dataset, identifies the community with the fastgreedy algorithm mentioned above, and records the propensity for each pair of individuals to be allocated to the same community. To test the robustness of non-breeding groups, we replaced the community detection process with the same assignment pairs of individuals that were observed together more than 75% of the time to the same group, and recorded the propensity for each pair of individuals to be allocated to the same group as in the original dataset. We ran the robustness algorithm 1000 times for each network, and evaluated how structured the population was in each of the seasons using the test statistic r_com_, representing the robustness of the assignment of individuals into the same community across all bootstrapped networks. An r_com_ value exceeding 0.5 suggests that the network is highly structured by communities and, therefore, that there are consistent preferences among individuals to associate (Shizuka & Farine 2016).

To confirm that non-breeding groups and communities in the networks were more strongly connected than expected by chance, we compared the mean association strength within groups and within communities of the real networks with those from 1000 permuted networks, using established pre-network permutations that swap pairs of 1s and 0s in the group-by-individual matrix (Farine 2017; Farine and Carter 2020).

#### Testing for community stability across years

We tested if the social composition of the communities was consistent across two consecutive non-breeding seasons. To do so, we first created a matrix of community co-membership for our two non-breeding season datasets. Given that the identity of the breeding groups was maintained throughout the study period (represented by high edge weights between individuals of the same breeding groups in network 2019: mean=0.93, SD=0.12 and network 2020: mean=0.90, SD= 0.11, see results), and that these could easily drive a high between-year correlation, we collapsed the breeding units into one node for this analysis. Thus, we generated a network as a matrix in which the rows and columns represented breeding units, and cell values given a 0 and 1 depending on whether two breeding groups were part of the same non-breeding community or not. We used a Mantel test to compare the breeding unit association matrix of the 2019 network with the one developed for the 2020 dataset to determine their similarity. High correlation values suggest consistent preferences among social units to associates across years.

#### Effect of home range overlap and breeding territory proximity on network structure

To determine the role of home range overlap in driving community structure, we calculated the kernel density estimation for the 95% of the home range of each group during the two years of study using the hadehabitatHR R package (Calenge 2015). For each non-breeding season, we created a network matrix of home range overlap by measuring the home range overlap of each group with all the others using the software QGIS (3.12). We then performed Mantel tests to quantify the correlations between networks representing group home range overlap and individual social networks.

To test if spatial proximity between breeding territories could influence social preferences and spatial overlap between individuals during the following non-breeding season, we calculated the position of the centroid for each breeding territory using the software QGIS (3.12). We then measured the distances between the centroids in all the breeding territories for breeding seasons (September-January) 2018-2019 and 2019-2020 using the R package Raster (Hijmans 2020), thus generating a matrix of spatial distances between breeding territories for each of the two breeding seasons. We ran Mantel tests to quantify the correlations between these spatial matrices of breeding territory centroids and those representing social connection between groups during the following non-breeding season.

## RESULTS

### Comparative analysis

We found sufficient data on presence/absence of potential MLS and cooperative breeding for 74 species out of the 113 species included in our research (see Table S2), with 22 species having evidence for a potential MLS (17 of 35 cooperative breeders and 5 of 39 non-cooperative breeders). For the remaining 38 species there was not enough information available on social organisation to classify the potential presence or absence of MLS. The comparative analysis indicated that cooperative breeders were more likely to be described using terms suggesting that they have a propensity to exhibit a MLS compared to non-cooperative breeders (46.1 %, *C.I*=25.95-67.67% probability vs 8.6 %, *C.I*= 4.97-14.35% probability; ß_non-cooperative breeders_= −2.21, SE= 0.75, p=0.003) (see Table S1). The amount of text devoted to the social organisation in the HANZAB did not significantly influence our classification of MLS (ß=0.14, SE=0.27, p=0.607). Full model output can be found in Table S1.

### Field case study

#### Social networks and multi-tier social structure in the superb fairy-wren

Our dataset comprised 1,632 social associations across two years, involving 113 individuals (60 males, 53 females) from 19 breeding groups. From the social networks, we identified three different stable social tiers. First, the upper-most mode of the distribution of association strengths suggested that this was comprised of individuals from the same breeding group/pair (units that are maintained during the breeding season) as well as some associations containing individuals from different breeding groups, and we called these associations between breeding groups/pairs supergroups. We then found that these breeding- and super-groups were embedded within robust and stable communities (sets of breeding groups/pairs that associate preferentially during the non-breeding season).

#### Non-breeding groups are composed of breeding groups and supergroups

From the two non-breeding season social networks, we identified groups as distinct clusters of individuals exhibiting very strong social bonds with each other (individuals found together more than 75% of the time) (see Fig.2). Robustness analysis of the co-membership of individuals in these groups confirmed that the study population had a clear group structure (non-breeding season 2019: r_com_=1; non-breeding season 2020 r_com_=0.98). Further, our analysis on the structure of non-breeding groups in the social networks against 1000 permuted networks indicated that these were significantly more strongly connected than expected by chance (p<0.001 for both years).

**Fig 1.**
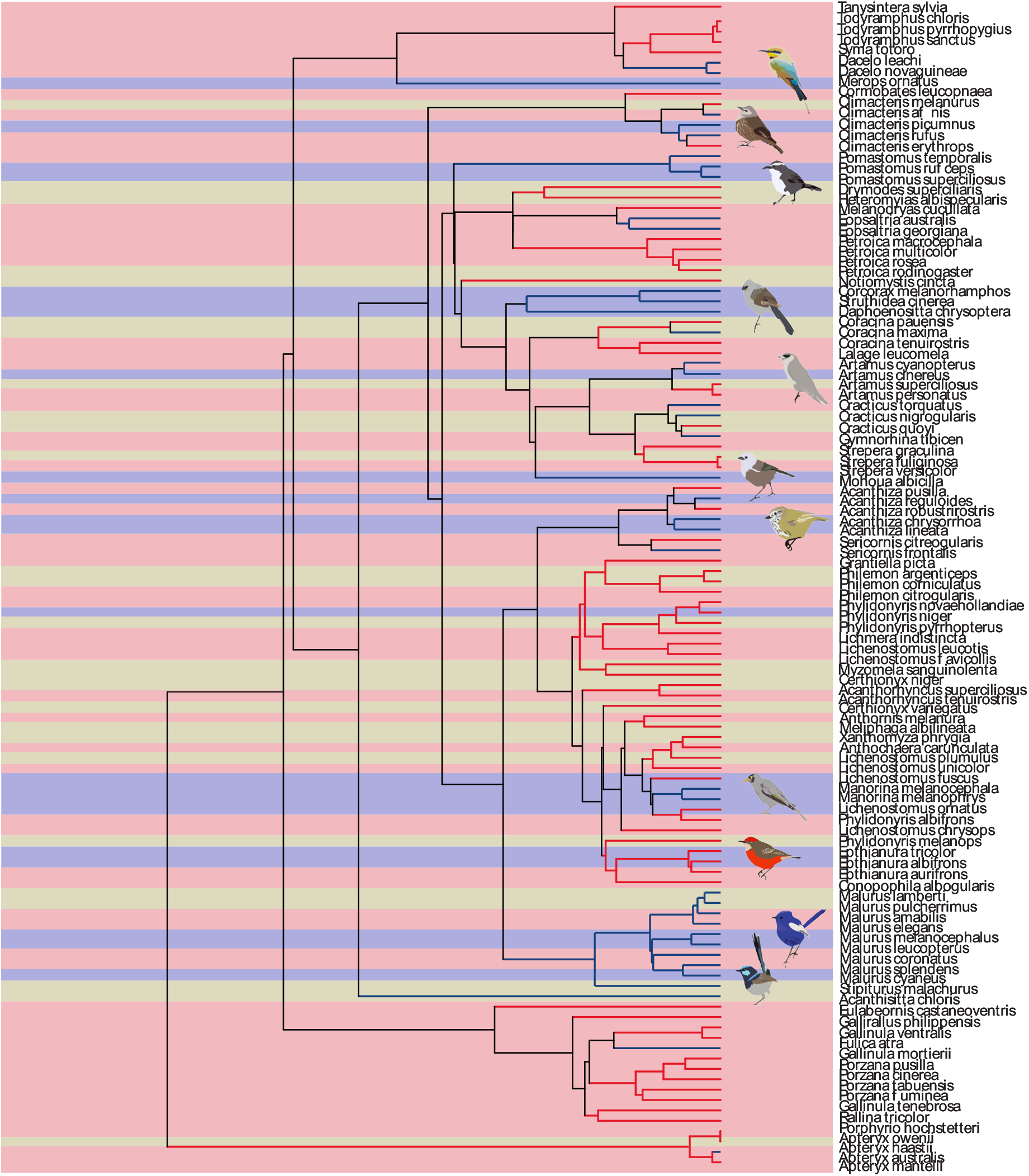
Overview of the phylogenetic relationships among Australian-New Zealand breeding birds. Colours of phylogenetic tree branches represent the presence (blue) or absence (red) of cooperative breeding. Blue background colour represents the probable existence of MLS, red represents the probable absence of MLS, while in yellow indicates the species for which we could not find enough evidence to suggest the presence or absence of MLS.

**Fig. 2.**
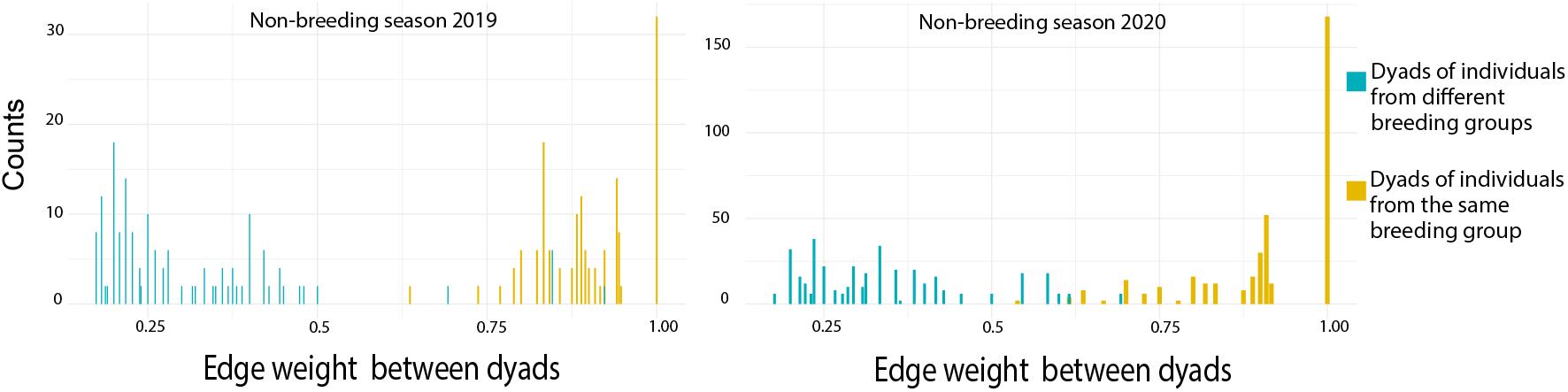
Base social tier of the MLS of superb fairy-wrens: individuals from the same breeding groups spend the non-breeding season together. Depicted is the strength of the social bond (edge weights) between dyads of individuals from 2019 and 2020 non-breeding season social network. Values above 0.75 show dyads forming part of the same non-breeding group, since they are consistently found together. Individuals that shared the same breeding groups the previous breeding season tend to maintain these social bonds during the non-breeding season by forming part of the same non-breeding group. Only edge weights higher than 0.17 are represented in the image.

In 2019, 11 of 15 groups were composed solely of members of the previous season breeding group, but four were formed by the members of two neighbouring breeding groups, forming supergroups (Fig.3). The same pattern was repeated in 2020, where 15 of the 17 clusters were composed of member of the previous season breeding group, while two formed supergroups (Fig.3). Moreover, the supergroups identified in 2020 were composed of the same breeding groups that had merged into supergroups in 2019, with those not re-observed also forming strong connections, highlighting their stability across years. A post-hoc analysis found that breeding group size was not related to supergroup formation (ß=0.14, SE=0.27, p=0.607). Rather, from the four supergroups observed during the course of the study, in at least three cases the dominant males of the two merging breeding groups had previously been part of the same breeding group. In all cases, one of the two dominant males was a previous subordinate of the other dominant male in the supergroup but later obtained its own breeding territory close to the natal territory. Interestingly, however, the converse was not always true, since two dominant males, which were previous subordinates near the present territory, did not join a supergroup. Unfortunately, we lack the data for the fourth supergroup since the dominant males were banded at the onset of the study in 2015, while they already occupied a dominant position.

**Fig 3.**
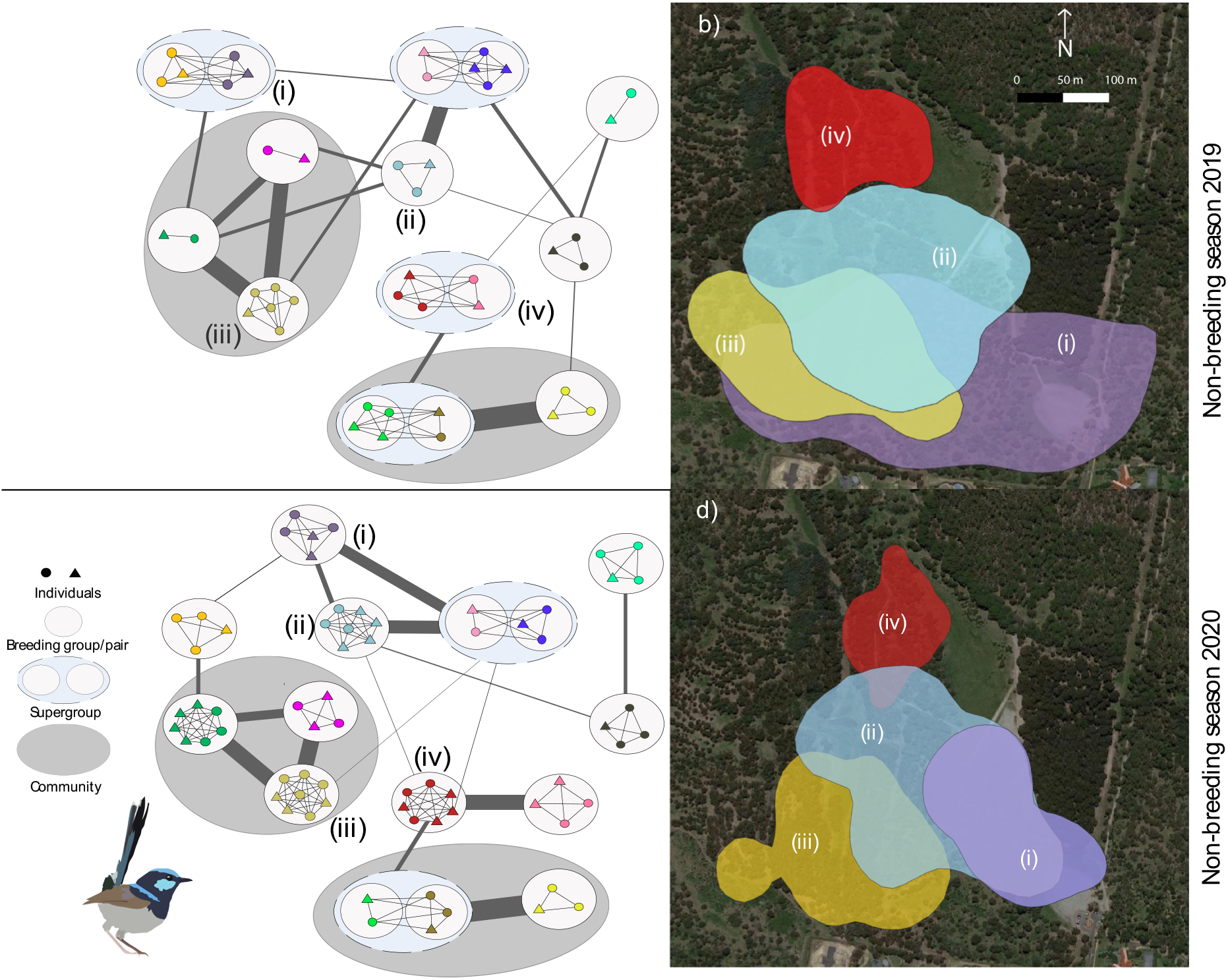
Overview of the multilevel structure of superb fairy-wren societies during the non-breeding season. Depicted is a portion of the network representing intergroup contacts in (a-b) 2019 and (c-d) 2020 non-breeding seasons. (a,c) At the lowest level of the society, colours refer to breeding group/pair membership with circles representing individual males and triangles representing individual females. Light grey ovals show non-breeding groups that are formed by breeding groups, while light blue ellipses with dashed line show non-breeding groups that correspond to supergroups. Dark grey ellipses represent examples of communities—sets of non-breeding groups that fission-fusion in and out of flocks, but have preferential associations with one-another. However, given the limited available space, only two communities per non-breeding season could be highlighted in the figure. Lines (in dark grey) connecting non-breeding groups represent whether they were observed together, with the thickness of the line showing the proportion of time each pair of non-breeding groups was in contact (highlighting that non-breeding flocks are not exclusive, but contain preferred associations). (b) The home range of four non-breeding groups represented by breeding groups (ii, iii) and two supergroups (i,iv) during the non-breeding season of 2019 and (d) the home range of the four non-breeding groups formed by breeding groups during 2020. The supergroups indicated by (i) and (iv) in 2019 split back into breeding groups during 2020 (d).

#### Communities

In both non-breeding networks, the fast-greedy statistical community-detection algorithm identified eight social communities in both the social networks. Analysis of the co-membership of individuals in these communities confirmed that the study population had clear community structure (2019: r_com_=0.79, 2020: r_com_=0.85), confirming the presence of preferential between-unit associations that drive the highest social level in fairy-wren societies (Fig. 3), while our analysis on the structure of communities in the social networks against 1000 permuted networks showed that these were significantly more strongly connected than expected by chance (p<0.001 for both years). Further, community social composition was correlated across the two years of study (Mantel test: *r*=0.53, *C.I.*=0.37-0.69, p<0.001), confirming that communities represent long-term social preferences that are generally maintained across non-breeding seasons.

#### Effect of home range overlap and breeding territory proximity on network structure

Distance between breeding territories during the breeding season significantly correlated with social structure during the non-breeding season. Mantel tests show that breeding groups with a larger breeding territory spatial distance had weaker social connection in non-breeding flocks [*r*= −0.66, *C.I.*=(−0.53,−0.79) for 2019 and *r*= −0.6, *C.I.*=(−0.45, −0.75), for 2020] and smaller home range overlap during the non-breeding season [*r*=−0.42, *C.I.*=(−0.25,−0.59) for 2019 and *r*=−0.7, *C.I.*=(−0.83,−0.57), for 2020], with all four correlations being statistically significant (at p<0.01). Mean spatial distance (±SD) between breeding territories of individuals being part of the same non-breeding community was 126±53 m in 2020 and in 2019 mean=122±73 m. Breeding group home ranges during the non-breeding season were significantly larger than their breeding territory size (2019 median size breeding group home range during non-breeding season=8.4 ha, IQR (Interquartile range, a measure equal to the difference between 75th and 25th percentiles)=5.6 ha; 2019 median size of breeding territory =0.39 ha, IQR=0.23; 2020 median size breeding group home range during non-breeding season = 5.0 ha, IQR=4.2 ha; 2020 median size of breeding territory=0.76 ha, IQR=0.39 ha). Home range size during the non-breeding season for each group varied between years (Wilcoxon test=260, p=0.02), as did the distance between breeding territory centroid and non-breeding home range centroid (2019 median distance =84 m, IQR=73 m, 2020 median distance=48 m, IQR=41 m) and a post-hoc analysis found that breeding group size did not predict home-range size (ß=1.16, SE=1.10, t=1.21).Many breeding groups showed large home range overlap during non-breeding season, with an average (±SD) overlap between groups home range of 36±30% during 2019 and 41±37% during 2020.

## DISCUSSION

Our comparative analysis of MLSs in cooperatively and non-cooperatively breeding birds shows that the emergence of MLSs in birds is likely facilitated by cooperative breeding and/or by similar factors to those that drive cooperative breeding. Our field case study on a cooperatively breeding species, combining year-round detailed observations of individually marked fairy-wrens with social network analyses, then revealed that superb fairy-wrens live in a highly-structured society. This society includes a multi-tiered structure outside of the breeding season, which is consistent between years. Together, these results support recent suggestions that MLSs are likely to be widespread in birds (Papageorgiou & Farine, 2020).

In superb fairy-wrens, three distinct, stable, hierarchical social units emerge over the year to form a multilevel society, with the social units at each level having stable membership within and across years. During the breeding season superb fairy-wrens live in exclusive territorial breeding groups. However, the boundaries between groups (and their territories) collapse after the breeding season. During the non-breeding season, breeding groups interact with other breeding groups in a predictable way, sometimes forming supergroups (the intermediate social tier; Fig. 3) and, in turn, merging to form communities (the upper social tier; Fig. 3). These communities are clusters of individuals that are tightly connected within the social networks of non-breeding flocks (Shizuka et al. 2014). During the following breeding season, communities and supergroups split back into breeding groups, and then reassemble again to re-form the same supergroups and communities after the subsequent breeding season.

The much larger home ranges during non-breeding compared to breeding home ranges (territory size) and the observed variability in home-ranges between years suggest that MLSs allow groups to exploit larger areas during the harsher season (winter) or years. This last argument is anecdotally supported by our data, where groups exhibited larger non-breeding home-ranges during the non-breeding season of 2019, which was preceded by four months (January-April) with dramatically less rainfall and higher maximum temperatures (harsher weather) compared to the same period in 2020 (see Supp. material 1). Future research could explore whether superb fairy-wrens respond to impacts of spring and summer rainfall on winter food availability (Cockburn et al. 2008) by adjusting their home range. Finally, an increase in predator pressure for adult individuals during the non-breeding season (unpubl. data) might increase the benefits of forming upper social units as a means of providing safety when foraging is more challenging (Kenward 1978; Delm 1990). Previous work in our study population found that fairy-wrens spend less time on vigilance behaviour and more time foraging when in larger groups (McQueen et al. 2017). However, forming larger groups is also risky. For example, it can increase female conflict over territories, breeding positions or partners. As a result, in species with high social viscosity, such as superb fairy-wrens, higher-level associations are likely to be preferentially formed among kin with established territories as a means of avoiding elevated levels of inter-group aggression.

We found that an intermediate social level (supergroup) can form from two breeding groups coming together, where dominant males were previously part of the same breeding group. There is a high probability of these males being related, given the pattern of positive spatial genetic correlation among males (radiating up to 160m in a different population; Double et al. 2005) and superb fairy-wren breeding groups within communities only rarely (14 %) occupying breeding territories located more than 160m apart. In fact, spatial distance between breeding territories strongly explained the strength of the social bonds between breeding groups and, thus, the social composition of communities (upper social level). We cannot distinguish whether the formation of communities in the superb fairy-wren is driven exclusively by active social preference or involves ecologically driven mechanisms (Farine 2017, He et al. 2019), but we hypothesise that social viscosity leading to kin-neighbourhoods (Hatchwell 2009) is likely to play a fundamental role in driving the emergence of upper social units (supergroups and communities) for the reasons described above.

Social viscosity is a key characteristic of cooperatively breeding birds, and our quantification of social structure in superb fairy-wrens formally captures what our literature review suggests is a widespread social behaviour among cooperatively breeding birds species across Australia and New Zealand. Our comparative analyses revealed that MLSs appear to be more commonly expressed by cooperatively breeding species. Descriptions of the social structure of at least 17 Australian-New Zealand cooperatively breeding species highlight a distinct social structure in which two or more stable social units seasonally emerge throughout the year, as a consequence of the associations of lower-level social units, whereas only five species of non-cooperatively breeding birds met the criteria to suggest they potentially live in a multi-tiered society (see Table 1 and Table S2). Furthermore, the occurrence of MLS varies among closely related cooperative breeders (Fig. 1). For example, several fairy-wren (*Malurus spp*) and treecreeper (*Climacteris spp*) species (i.e. purple-crowned fairy-wren, splendid fairy-wren and white-browed treecreeper) have strict year-round territoriality which, inherently, cannot exhibit multi-tiered social structure. Yet, these two genera also include species showing potential multilevel social system, where kin breeding groups interact to form higher social units outside the breeding season (i.e. superb fairy-wren and white-winged fairy-wren) or during the breeding season (brown treecreeper). Among the five non-cooperatively breeding species potentially exhibiting MLS, chats and white-cheeked honeyeater appear to form stable flocks (higher social unit) during the non-breeding season which can then split back into pairs of individuals defending territories during breeding season, a similar pattern to superb fairy-wrens and other cooperatively breeding species (see Table 1). Yellow-plumed honeyeater and fuscous honeyeater are semi-colonial species, reported to form well-defined clusters of individuals that can associate to mob intruders (see Table 1). Their social structure is reminiscent of that found in miners, cooperative breeders which have clearly defined layers of social organisation. However, further studies are necessary in order to better explore the extent of the similarities and differences in the societies of these species.

**Table 1.**
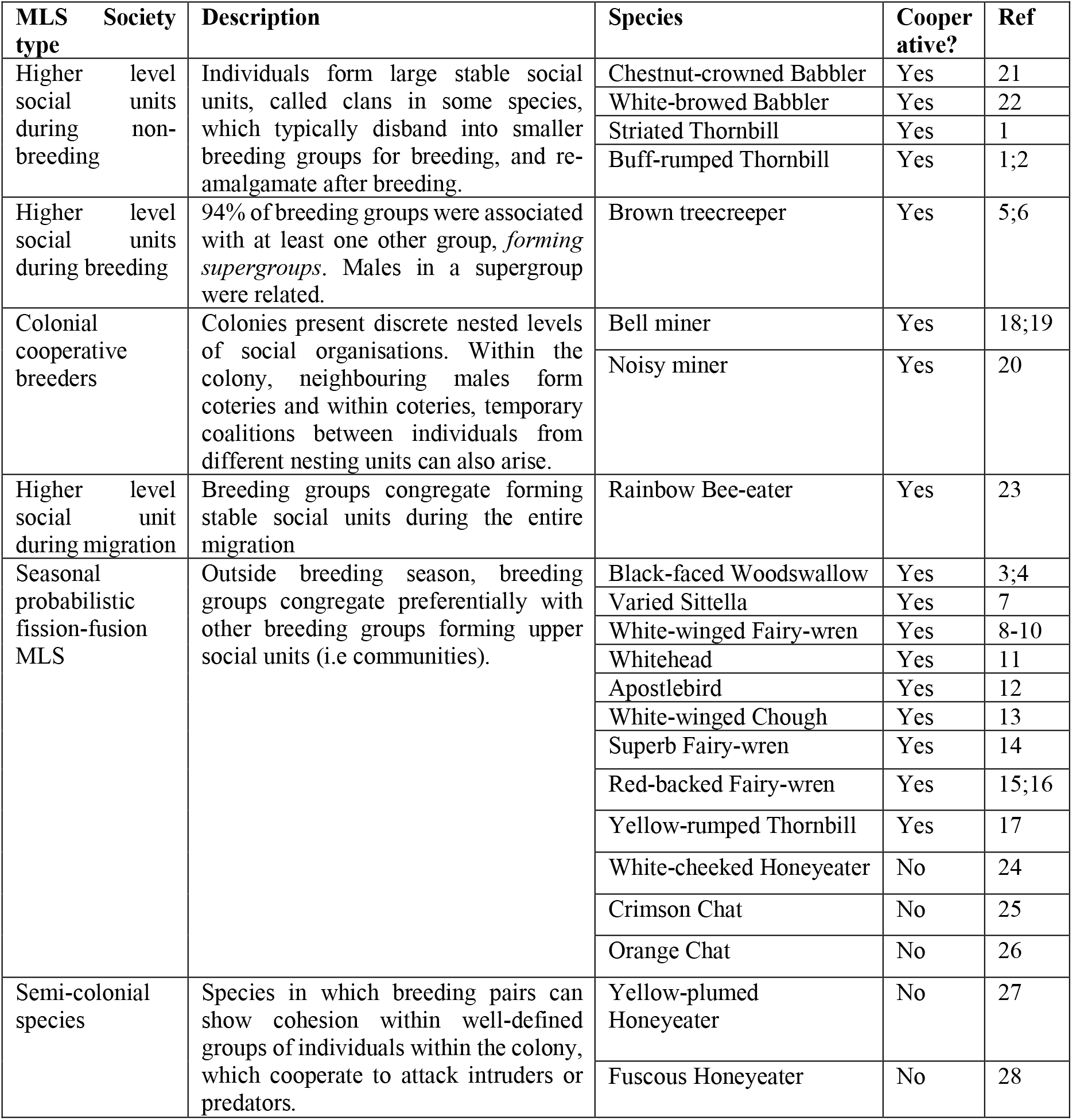
Description of putative types of MLS across bird species from Australia and New Zealand. We found support for a strong indication of multilevel society structure in 17 of 35 cooperatively breeding species, but only five of 39 non-cooperative species included in our search.

The weak phylogenetic signal in the probability of multilevel social organization (see Table S1) suggests that it is a relatively fluid social trait that can emerge and disappear, and highlights how birds can offer new opportunities to identify drivers that lead to the emergence of MLS (Papageorgiou and Farine 2020). For example, some species, such as cooperatively breeding yellow-rumped thornbills for which we did not find evidence to suggest they form MLSs, form non-breeding mixed-species flocks with those that do (such as buff-rumped thornbills; Farine and Milburn 2013). During the non-breeding season, superb fairy-wrens also regularly form mixed species flock with other passerine species, some of which are likely to exhibit MLS (i.e striated thornbills; Bell 1980). This suggests not only that yellow-rumped thornbills could express a multilevel-like social structure involving members of a different species, but also raises questions about how mixed-species associations between cooperatively breeding species and other species (e.g. *Petroicidae* robins, *Climacteridae* treecreepers, and many of the *Acanthizidae* species including other thornbills; see Bell et al. 1980) fit into the MLS framework, especially given that many of these associations may not be random at the individual level (Farine and Milburn 2013). In any case, the formation of multi-species MLS cannot be driven by family-ties, and therefore comparisons of single- and multi-species MLS could provide insights into the relative importance of different mechanisms and factors that shape MLS (like spatial distribution of resources or reduced predation risk).

There are striking similarities between the non-breeding social structure of cooperatively breeding superb fairy-wrens and a phylogenetically distant species that also has an MLS, the vulturine guineafowl. In both societies, for example, the groups show a biased sex-ratio and are centred around the philopatric sex, groups significantly expand their home ranges during harsher seasons (Papageorgiou et al. 2021), and it is likely that preferred associations among guineafowl groups are also linked to relatedness among males (D.R. Farine, *pers. obs*.) due to a strong female bias in dispersal (Klarevas-Irby et al. 2021). The high social viscosity that emerges from limited dispersal in cooperatively breeding species might then provide the foundations for the formation of upper social units (Bell and Ford 1986; Painter et al. 2000). Relatedness also appears to play an important role in mammalian MLSs. For example, as the level of associations decreases and the strength of association increases (i.e. from community, to band, to team, to unit) in geladas (*Theropithecus gelada*), females are more likely to be genetically related (Tinsley Johnson et al. 2014). Yet, to our knowledge, none of the well-known mammalian MLSs are also cooperative breeders. This raises the question whether different mechanisms ‘set the scene’ for the emergence of MLSs (social viscosity) among birds and mammals, even if the drivers underlying the formation of MLSs (e.g. environmental harshness, the benefits of grouping, and avoiding conflict with non-relatives) are similar.

One of the main open questions in the study of social evolution is what conditions and ecological pressures favour the emergence of MLS (Grueter et al. 2020). The proximate mechanisms underlying the formation of MLSs are presumed to vary widely (Grueter et al 2020), and expanding the focus of research to other taxa can enhance the insight we can gain on the drivers of complex social structure (Papageorgiou and Farine, 2020). We encourage researchers who study social behaviour of cooperatively breeding species to investigate the social structure of their species during the non-breeding season, which will help to identify generalities and exceptions to the pattern we uncovered here. Another promising research avenue is the study of the non-breeding social behaviour of the non-cooperative species highlighted in our study as potentially displaying MLS, which might further enhance our understanding of the ecological drivers of MLS. Explicitly including the often neglected non-breeding period in studies of social behaviour could offer deeper insight into the social complexity of these species, and a better understanding of the drivers of MLS across animals.

## Supporting information

Supplemen Camerlenghi et al.

## ACKNOWLEDGEMENTS

We thank Catherine Villeneuve, Sergio Nolazco Plasier, Jenna Diehl, Abigail Robinson, Gregory Taylor, Niki Teunissen and many volunteers for their help in the field. We thank three anonymous reviewers for their comments which improved the manuscript. Fieldwork was approved by the Animal Ethics Committee of the School of Biological Sciences of Monash University (BSCI/2016/03 and 16348) and approved by Department of Environment, Land, Water and Planning and Parks Victoria (permit no. 10008307 and 10008704) and the Australian Bird and Bat Banding Scheme (ABBBS nr. 2230). Funding was provided by the Holsworth Wildlife Research Endowment and the Ecological Society of Australia (to EC), the Australian Research Council (DP180100058 to AP) and Monash University. DRF was funded by the European Research Council (ERC) under the European Union’s Horizon 2020 research and innovation programme (grant agreement number 850859) and an Eccellenza Professorship Grant of the Swiss National Science Foundation (grant number PCEFP3_187058).

## Table 1 references

[1] HANZAB vol 6, pag 541. [2] HANZAB vol 6, pag 500. [3] HANZAB vol 7a, pag 445. [4] Rowley 1999. [5] HANZAB vol 5, pag 232. [6] Doerr, E. and Doerr, V.A.J. (2006). *Anim Behav*. [7] HANZAB vol 6, pag 1007. [8] Rowley and Russel. (1995). *Emu*. [9] Rathburn, M.K. and Montgomerie R. (2003). *Emu*. [10] HANZAB Vol 5, pag 352. [11] HANZAB vol 6, pag 1027. [12] Griesser et al. (2009). *Ethology*. [13] Heinson, RG. (1987). *Behav. Ecol. Sociobiol*. [14] Rowley, I. (1964). *Emu*. [15] Rowley I and Russell E. (1997). *Fairy-Wrens and Grasswrens*. Pag 183; [16] Macgillivray, W. (1914). *Emu*. [17] HANZAB vol 6, pag 510. [18] Bell, H.L. and Ford, H.A. (1986). *Behav. Ecol. Sociobiol*. [18] HANZAB vol 5, pag 615; [19] Painter at al. (2000). *Mol. Ecol*. [20] HANZAB vol 5, pag 631-632. [23] Garnett, S. (1985). *Emu*. [24] HANZAB vol 5, pag 1044-45. [25] HANZAB vol 5, pag 1212-1214. [26] HANZA vol 5, pag 1203-1204. [27] HANZAB vol 5, pag 843-844. [28] HANZAB vol 5, pag 863.

## Notes

### Competing Interest Statement

The authors have declared no competing interest.

