## Supplementary material for "Cooperative breeding and the emergence of multilevel societies in birds": Supplemen Camerlenghi et al.

### Supplement

During January-April 2019 the mean maximum temperature was 27.1°C (SD=6.5°C) and the total amount of rainfall was 106 mm. During the same months in 2020, the mean maximum temperature was 23.4°C (SD=5.4°C) and the total amount of rainfall was 533 mm. During winter months (May-August) 2019, the mean minimum temperature was 7.3°C (SD=3.1°C) and the total amount of rainfall was 337 mm. During the same months in 2020, the mean minimum temperature recorded was 6.4°C (SD=3.3°C) and the total amount of rainfall was 257 mm.

Data was recorded by the Scoresby research institute (-37.8710 145.2561), situated at 8.8 km from the study site, and it was freely accessible at the Bureau of Meteorology of the Australian Government web-page (http://www.bom.gov.au/climate/data/).

**Table S1.** Results of phylogenetic logistic regression analysing the effects of cooperative breeding (yes/no) and amount of knowledge (the length of text devoted to the social organisation in Marchant et al. 2006) on the presence/absence of MLS in Australian birds belonging to families that have at least one species showing cooperative breeding. Depicted are average effects and associated statistics across 100 phylogenies to account for phylogenetic uncertainty. Model alpha (an indicator of phylogenetic signal, where lower values correspond to stronger phylogenetic effects) was 0.134, indicating a relatively weak phylogenetic signal.

| **effect** | **estimate** | SE | **t-value** | **p-value** |
|---|---|---|---|---|
| Intercept | -0.155 | 0.456 | -0.341 | 0.732 |
| Non-coop. vs cooperative | -2.214 | 0.753 | -2.937 | 0.003 |
| Text length HANZAB | 0.140 | 0.273 | 0.514 | 0.607 |
